# Cushing’s Syndrome mutant PKA^L205R^ exhibits altered substrate specificity

**DOI:** 10.1101/091231

**Authors:** Joshua M. Lubner, Kimberly L. Dodge-Kafka, Cathrine R. Carlson, George M. Church, Michael F. Chou, Daniel Schwartz

## Abstract

The PKA^L205R^ hotspot mutation has been implicated in Cushing’s Syndrome through hyperactive gain-of-function PKA signaling, however its influence on substrate specificity has not been investigated. Here, we employ the Proteomic Peptide Library (ProPeL) approach to create high-resolution models for PKA^WT^ and PKA^L205R^ substrate specificity. We reveal that the L205R mutation reduces canonical hydrophobic preference at the substrate P+1 position, and increases acidic preference in downstream positions. Using these models, we designed peptide substrates that exhibit altered selectivity for specific PKA variants, and demonstrate the feasibility of selective PKA^L205R^ loss-of-function signaling. Through these results, we suggest that substrate rewiring may contribute to Cushing’s Syndrome disease etiology, and introduce a powerful new paradigm for investigating mutation-induced kinase substrate rewiring in human disease.

## Introduction

Cushing’s Syndrome is defined by a collection of signs and symptoms that result from prolonged hypercortisolism (due to excessive secretion of adrenocorticotropin hormone, or adrenocortical adenomas), with patients commonly presenting with central obesity, metabolic anomalies, and hypertension [1]. At a biochemical level, cAMP-dependent protein kinase A (PKA) has been implicated in Cushing’s Syndrome through several different lines of evidence, and PKA mutations have been identified in almost 40% of cortisol-secreting adenomas [2]. Like other protein kinases, PKA derives substrate specificity through temporal and spatial co-localization, scaffolding, and docking interactions. Kinase specificity is also determined by interactions with specific patterns of amino acids surrounding the phosphoacceptor, which is referred to as the “kinase specificity motif” [3]. Recently, several groups genetically identified the hotspot mutation PKA^L205R^ in patients diagnosed with Cushing’s Syndrome [4–7]. In PKA^WT^, residue L205 (together with residues L198 and P202) form what is known as the P+1 loop, where P+n denotes the nth residue towards the C-terminus of the phosphoacceptor (P-site), and P-n denotes the nth residue towards the N-terminus. The P+1 loop contributes to the PKA^WT^ specificity motif by creating a hydrophobic pocket that favors substrates with a P+1 hydrophobic residue [8,9].

Residue L205 is highly conserved in PKA homologs from humans to invertebrates, and also plays an important role in stabilizing the hydrophobic interaction with I98 in the regulatory subunit RIα pseudo-substrate sequence RRGAI (and the homologous positions in regulatory subunits RIβ, RIIα and RIIβ [10]). The role of L205 in forming the P+1 loop and an observed increase in cAMP signaling associated with the PKA^L205R^ mutant has led to the prevailing hypothesis that mutation from leucine to arginine disrupts the interface with regulatory subunits, resulting in a constitutively active PKA enzyme [11].

Recently, kinase missense mutations identified from cancer patients were shown to modulate not only catalytic activity, but in some cases to alter substrate specificity [12]. Until now, there has been no investigation into the effect of the L205R mutation on PKA substrate specificity. The structures of PKA^WT^ and PKA^L205R^ in complex with the naturally-occurring inhibitor peptide PKI (containing the pseudo-substrate sequence RRNAI) reveal a distinct reorientation of the P+1 isoleucine in PKI when in complex with PKA^L205R^, with no other apparent changes in kinase-substrate binding (Fig. 1A and Fig. S1, [13]). This change in binding is likely to accommodate the loss of the P+1 loop hydrophobic pocket in the mutant, due to introduction of hydrophilic character and steric clash with the bulky arginine side-chain. Based on this structural data, we hypothesized that PKA^L205R^ would exhibit a reduced preference for substrates with a hydrophobic residue in the P+1 position, while maintaining canonical upstream basophilic interactions.

**Figure 1.**
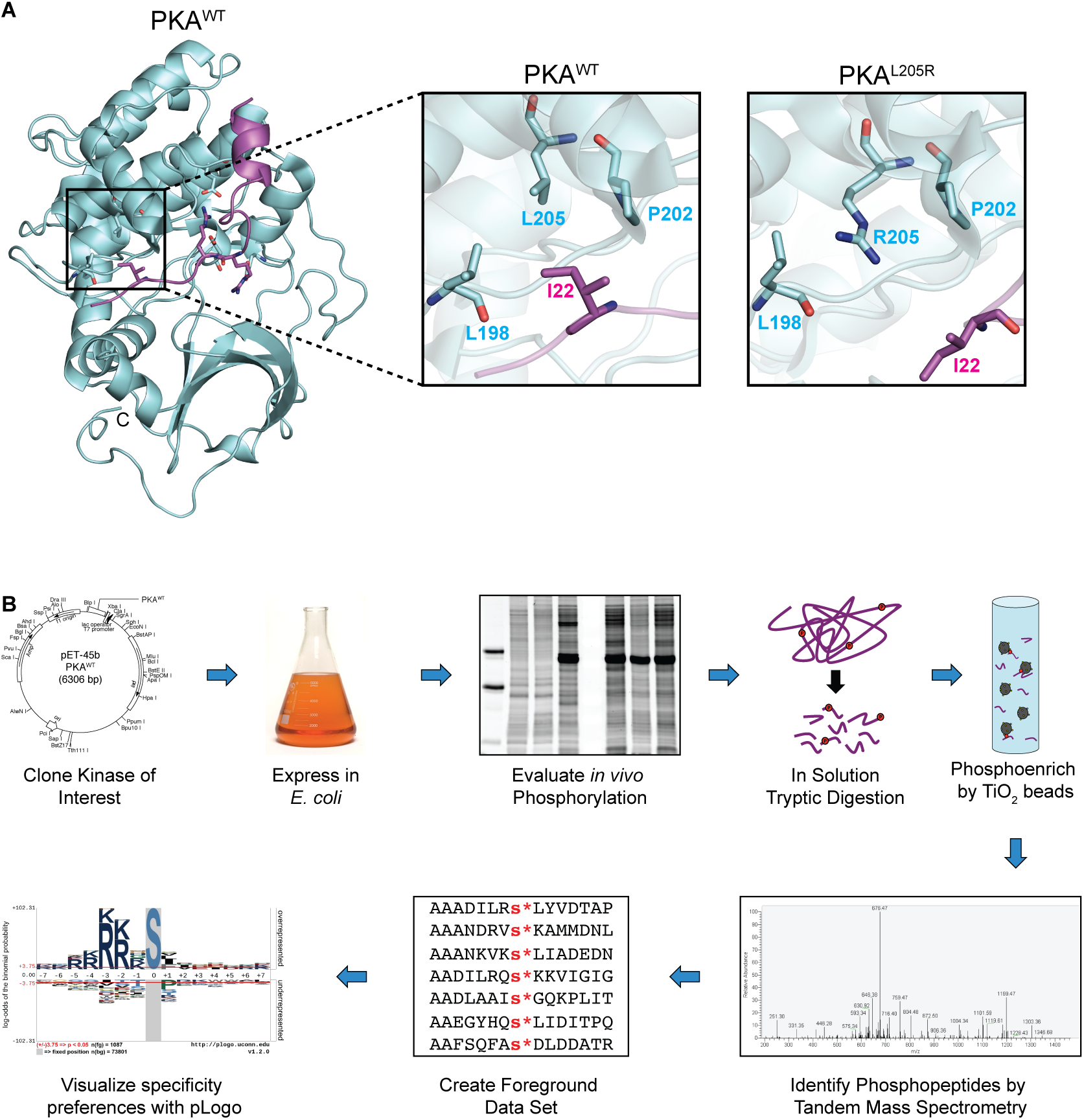
**Experimental rationale and workflow.** (A) Structural visualization of the substrate peptide binding site for PKA^WT^ (PDB Model 4WB5 [13]) indicating putative hydrophobic interaction between PKA residues L198/P202/L205 and PKI substrate I22 at P+1. In the Cushing’s syndrome mutant PKA^L205R^ (PDB Model 4WB6 [13]), this pocket has been destabilized. PKA is colored in cyan, with peptide substrate in magenta. Expanded views of both structures are available in Fig. S1. (B) Schematic overview of the experimental ProPeL workflow. A kinase of interest is cloned, and expressed in *E. coli*. Resulting bacterial phosphorylation is evaluated by SDS-PAGE with Pro-Q Diamond and coomassie staining. Lysate is digested, phosphoenriched and identified by tandem mass spectrometry. Data sets are visualized with pLogo [19].

## Materials and methods

### Plasmids, strains, and *in vivo* proteome phosphorylation

A plasmid containing the full-length coding sequences for the human PRKACA gene in the pDNR-Dual vector was purchased from the Harvard PlasmID Repository (Boston, MA). Full-length PKA was cloned into the pET45b vector (Novagen) by traditional restriction site PCR cloning. The PKA^L205R^ mutation was created according to the Stratagene QuickChange II protocol. PKA constructs were transformed into the *E. coli* Rosetta2 strain (Novagen) and expressed by IPTG induction. PKA expression was optimal when induced at mid-log with 0.5 mM IPTG and incubated overnight at 22°C with shaking at 250 rpm. Cells were harvested by centrifugation at 6,000 g and 4°C for 10 minutes, and stored at -80°C.

### Lysis and analysis of *in vivo* phosphorylation

Cell lysate was prepared as described previously [14,15] with minor modifications. Cells were lysed by sonication with a Fisher Sonic Dismembrator F60 at 15% power using 15–20 second pulses, with 1 minute rest on ice between pulses, until lysate was clear. Crude lysate was clarified by centrifugation at 20,000 g and 4°C for 30 minutes. Protein concentrations were determined by Bichinchoninic Acid (BCA) Assay (Pierce), phosphorylation level was evaluated by SDS-PAGE with Pro-Q Diamond Phosphoprotein stain (Life Technologies), and total protein was evaluated by GelCode Blue coomassie staining (Life Technologies).

### Western blotting

Western blotting for PKA used primary rabbit antibody for the human catalytic subunit α (PKAα cat (C-20): sc-903, Santa Cruz Biotechnology) at 1:1000 dilution, and IRDye 680RD Donkey anti-Rabbit IgG secondary antibody (LI-COR Biosciences) at 1:5000 dilution.

### In-solution tryptic digestion

Samples were reduced, alkylated, digested with trypsin (Promega) at a 1:100 enzyme:substrate ratio, and desalted as previously described in Villén and Gygi, steps 2-17 [14].

### TiO_2_ bead phosphopeptide enrichment

Phosphopeptide enrichment using bulk TiO_2_ beads (Titansphere 5 µm, GL Sciences) was modified from Kettenbach and Gerber [16]. Beads were conditioned in bulk using Binding Buffer (50% ACN, 2M Lactic Acid), with beads added at a 4:1 ratio to peptides (peptide concentration estimated by NanoDrop A280 absorbance), and brought to a final peptide concentration of 1 mg/mL. Peptide/bead mix was incubated with maximum shaking on an Eppendorf Thermomixer at room temperature for 1 hour. Beads were washed with Wash Buffer (50% ACN, 0.1% TFA) and eluted with 5% NH_4_OH. Eluate was immediately acidified by addition of FA, dried in a speed-vac, and stored at -20°C until analysis by mass spectrometry.

### Peptide identification by tandem mass spectrometry

Peptides were resuspended in 30 µL Buffer A (3% ACN, 0.125% FA) and 1-4 µL loaded onto a C18 nanocapillary column with a pulled tip that sprays directly into the inlet of a Thermo Fisher Scientific LTQ Orbitrap XL mass spectrometer. Peptides were eluted using an Agilent 1200 HPLC binary pump with a gradient that changes solvents from 100% to 65% Buffer A (0% to 35% Buffer B) over a 75 minute time period, where Buffer A = 3% ACN, 0.125% FA in water, and Buffer B = 0.125% FA in ACN. A TOP10 method was used (MS scans followed by Collision Induced Dissociation MS/MS on the top 10 most intense MS spectral peaks). Spectra were searched using SEQUEST against the *E. coli* proteome, including decoy database entries, which allowed for differential serine, threonine, and tyrosine phosphate modifications (+79.966331), a differential methionine oxidation modification (15.9949146221) and a constant cysteine modification of +57.02146374. The deltaXCORR (the difference between the first and second hits to the databases) was set to be >= 0.08. To minimize false positives, for each of the two classes of peptide charges z = +2 and z >= +3, XCORR thresholds were chosen to accept peptides in such a manner that 1% of them were hits from the decoy database, resulting in an expected False Discovery Rate (FDR) of 2%.

### Phosphopeptide list filtering

Prior to motif analysis, a master negative control list was generated by pooling phosphopeptides previously identified in negative control experiments [15], previously identified endogenous *E. coli* phosphorylation sites [17,18], and phosphorylation sites identified in empty vector and kinase dead negative control experiments. Phosphorylation sites on this master negative control list were removed from each PKA variant data set to generate a final list of kinase-specific phosphorylation sites. Peptide lists from all runs were merged within each kinase variant, and redundant peptides were removed prior to motif analysis.

### pLogo generation

To generate graphical motifs, known as pLogos, we used the online tool at plogo.uconn.edu, previously described in detail [19]. See Supporting Information for a more detailed explanation, and instructions for recreating pLogos with our provided data.

### Structural modeling

All structural modeling was visualized using PyMol for Mac, and using the following structures retrieved from the RCSB Protein Data Bank: PKA^WT^ – PDB Model 4WB5, and PKA^L205R^ – PDB Model 4WB6 [13].

### Recombinant kinase purification

PKA^WT^ and PKA^L205R^ were expressed as described above. Pelleted cells were resuspended in a native Ni-NTA A buffer (50 mM Tris-HCl, pH 8, 0.5 M NaCl, 20 mM Imidazole, 2 mM DTT, 10% glycerol) supplemented with Halt Protease Inhibitor Cocktail EDTA-free, Halt Phosphatase Inhibitor Cocktail (Pierce) and 1 mM phenylmethylsulfonyl fluoride, and then sonicated, clarified, and quantified by BCA assay as described above. For each PKA variant, 15 mg clarified lysate was incubated with 250 µL HisPur Ni-NTA resin (Thermo Fisher Scientific) and brought to a final volume of 1 mL with Ni-NTA A buffer. Slurry was incubated with end-over-end rotation at 4°C for 1 hour. Resin was washed by gravity flow on ice with 10 mL Ni-NTA A buffer, and eluted with 1.5 mL Ni-NTA B buffer (same as Ni-NTA A with 0.25 M Imidazole). Eluate was dialyzed against storage buffer (50 mM NaCl, 20 mM Tris-HCl, pH 7.5, 1 mM EDTA, 2 mM DTT, 25% glycerol), snap frozen in liquid nitrogen, and stored at -80°C.

### Peptide array membrane synthesis and [γ-^32^P]ATP assay

Peptides were synthesized as 7- or 15-mers on membranes using a MultiPep automated peptide synthesizer (INTAVIS Bioanalytical Instruments AG, Koeln, Germany). The kinase assay protocol was modified from Himpel et al. [20]. Membranes were blocked overnight in 15 mL kinase buffer (20 mM Hepes, pH 7.4, 5 mM MgCl_2_, 1 mM DTT) with 0.2 mg/mL BSA and 100 mM NaCl. Blocked membranes were incubated with 15 mL fresh kinase buffer with 1 mg/mL BSA, 100 mM NaCl and 50 µM cold ATP at 30°C for 45 minutes. Kinase assays were conducted by incubating the membranes in 15 mL of fresh kinase buffer containing 0.2 mg/mL BSA, 12.5 µCi [γ-^32^P]ATP, and 1 µg of each kinase for 30 minutes at 30°C with slight agitation. Membranes were washed with 15 mL 1 M NaCl for 5 min, for a total of 5 washes, followed by 3 water washes. Membranes were then washed with 15 mL 5% phosphoric acid for 15 min, for a total of 3 washes, followed by 3 water washes. Membranes were air dried, and exposed to film for 3 days. Phosphorylation was analyzed by densitometry using Gilles Carpentier's Dot-Blot-Analyzer macro for ImageJ (written by Gilles Carpentier, 2008, and available at http://rsb.info.nih.gov/ij/macros/toolsets/DotBlotAnalyzer.txt). See Table S3 for raw data, and normalization protocol.

## Results

### Human PKA^WT^ and PKA^L205R^ are active when expressed in *E. coli*

In order to accurately determine specificity motifs for both PKA^WT^ and PKA^L205R^, we used the <u>Pro</u>teomic <u>Pe</u>ptide <u>L</u>ibrary (ProPeL) method [15] because it allowed us to determine high-fidelity motifs for individual kinase clones expressed in isolation (i.e. in the absence of potentially confounding eukaryotic kinases). Using this approach (Fig. 1B), first a heterologous kinase of interest is expressed in *E. coli*. Upon expression, the kinase phosphorylates bacterial proteins consistent with its intrinsic kinase specificity motif. After cell lysis and proteolysis, the resulting *E. coli* phosphopeptides are identified by tandem mass spectrometry. This can provide hundreds to thousands of kinase-specific phosphopeptides from which a high-resolution motif model is generated, and easily visualized using the pLogo graphical representation [19].

As per the ProPeL method, we expressed the full-length human catalytic subunit for either PKA^WT^ or PKA^L205R^ in the Rosetta2 *E. coli* strain, noting robust expression as evaluated by western blot (Fig. S2A). Using the in-gel phosphoprotein stain Pro-Q Diamond [21], we observed that both PKA variants exhibit strong autophosphorylation and efficiently phosphorylate *E. coli* proteins throughout the gel (and thus across the proteome) compared to the empty vector control (Fig. S2B,C). Lysate from cells expressing either PKA^WT^ or PKA^L205R^ was digested, enriched, and subjected to phosphopeptide identification by tandem mass spectrometry. Combined with data from previously published results [15], we obtained 1,404 unique phosphorylation sites (1,087 pSer and 317 pThr) for PKA^WT^, and 585 unique phosphorylation sites for PKA^L205R^ (445 pSer and 140 pThr, see Table S1).

### PKA^L205R^ exhibits altered substrate specificity

PKA^WT^ has a well-established specificity motif that consists of upstream basic residues (particularly at the P-2 and P-3 positions) and a significant enrichment for the hydrophobic (ϕ) residues [/I/L/M/V/F] at P+1. This motif can be simplified to the consensus sequence [R/K][R/K]×Sϕ [9,22]. While the PKA^WT^ pLogo generated from our dataset faithfully recapitulates this known motif (Fig. 2A), the PKA^L205R^ pLogo reveals a striking loss of hydrophobic preference (with the exception of leucine) at the P+1 position (Fig. 2B). In addition, we observe a strong [I/V] hydrophobic grouping at P+2 position in the PKA^WT^ motif, which is replaced by the acidic [D/E] grouping at this position in the PKA^L205R^ motif. This shift towards an increased enrichment for downstream acidic residues in the motif may be due to an electrostatic interaction with a positively charged R205 guanidinium group in the mutant kinase. Conversely, we note that the canonical basophilic preference at substrate positions P-2 and P-3 by PKA^WT^ was unaltered for PKA^L205R^, which is consistent with conserved interactions with PKI in the mutant crystal structure (Fig. S1). PKA^L205R^ motifs for substrates that contain threonine P-sites are largely similar to the serine motifs, although there appears to be an even greater shift away from hydrophobic P+1 preference and a less pronounced increase in acidic enrichment at downstream positions (Fig. S3). Together, these results demonstrate that the Cushing’s syndrome mutant PKA^L205R^ does indeed alter PKA substrate specificity, and in a manner predicted by the existing crystal structure data. The processed ProPeL experimental data are provided in Table S2. Interested readers may follow the detailed instructions for re-creating these pLogos to dynamically explore the correlated positions within the motifs (see Supporting Information).

**Figure 2.**
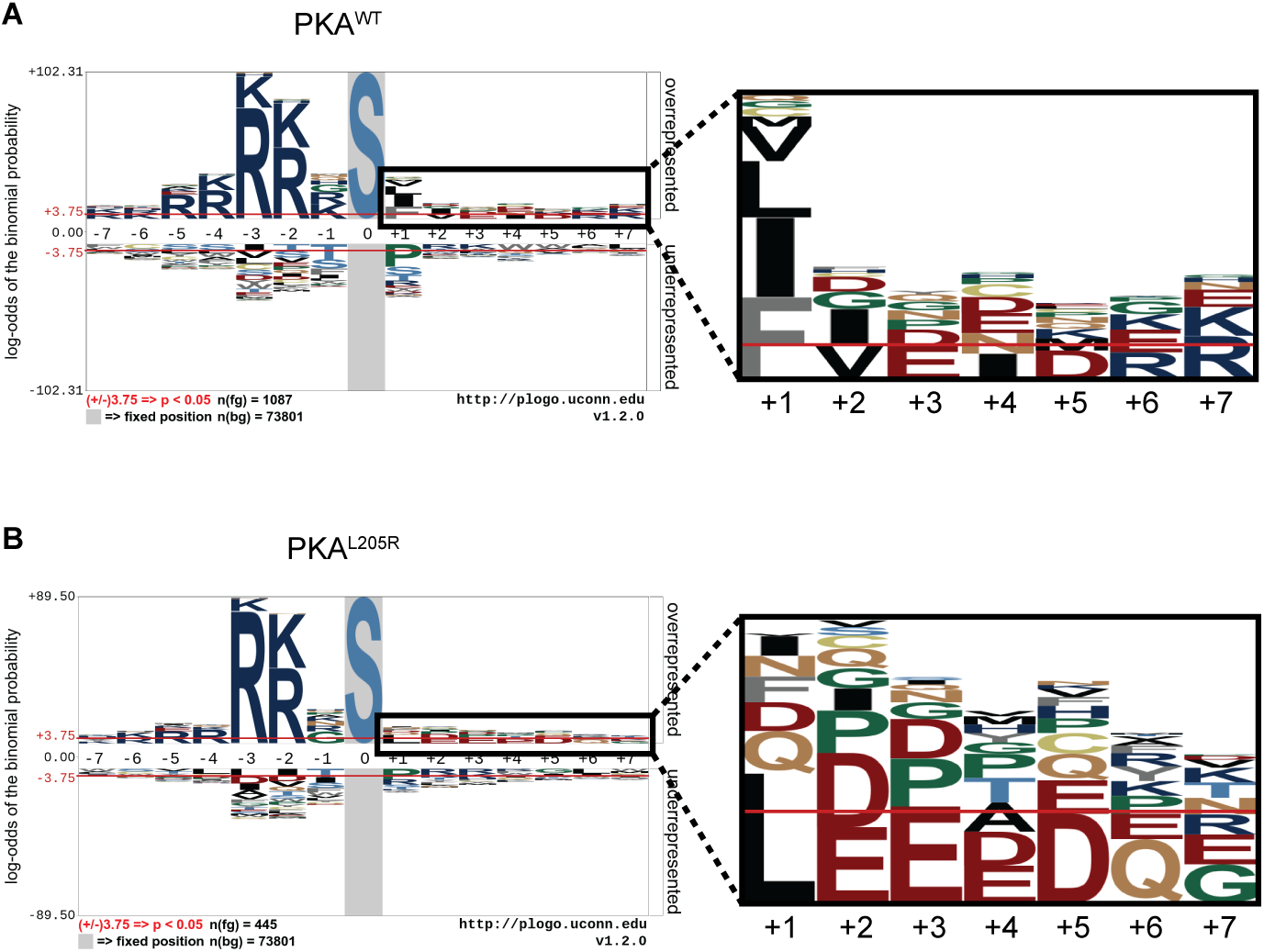
**PKA**^**L205R**^ **exhibits altered downstream substrate specificity.** (A to B) pLogos [19] illustrate substrate preferences for serine substrates of (A) PKA^WT^ and (B) PKA^L205R^. Zoom (A,B): zoomed in view of P+1 to P+7 positions. Overrepresented residues are displayed above the x-axis, underrepresented residues are below the x-axis. The *n(fg*) and *n(bg*) values at the bottom left of the pLogo indicate the number of aligned foreground and background sequences used to generate the image, respectively. The red horizontal bars correspond to *p* = 0.05 (corrected for multiple hypothesis testing), and y-axis is logarithmic scale. The grey box indicates a “fixed” residue. Additional pLogos for threonine-centered substrates are available in Fig. S3.

### Using pLogos as a guide for substrate engineering and prediction

To corroborate the results from our ProPeL-derived pLogos, we conducted [γ-^32^P]ATP *in vitro* kinase assays by incubating recombinant PKA with custom peptide arrays. First, we evaluated whether the observed *E. coli* peptide substrates identified by ProPeL were suitable PKA substrates *in vitro*. Using our PKA^WT^ pLogo and an internal version of the *scan-x* tool [23], we scored the *E. coli* phosphopeptides identified in the PKA^WT^ ProPeL experiments and selected two of the highest scoring peptides for synthesis (Observed Peptide 1 – PLRMRRGSIPALVNN, Observed Peptide 2 – IVLPRRLSDEVADRV). As a benchmark, we also selected a known human PKA^WT^ substrate for synthesis (CDK16^S119^ – EDINKRLSLPADIRL [24]). The *in vitro* kinase assay revealed that both the observed *E. coli* peptides and the human CDK16^S119^ peptide were readily phosphorylated (Fig. 3A), and a one-way ANOVA found that the means are not significantly different [*F*(2,14) = 0.56, *p* = 0.58349], verifying that phosphorylation sites obtained through ProPeL *E. coli* experiments are a suitable proxy for real human substrates.

**Figure 3.**
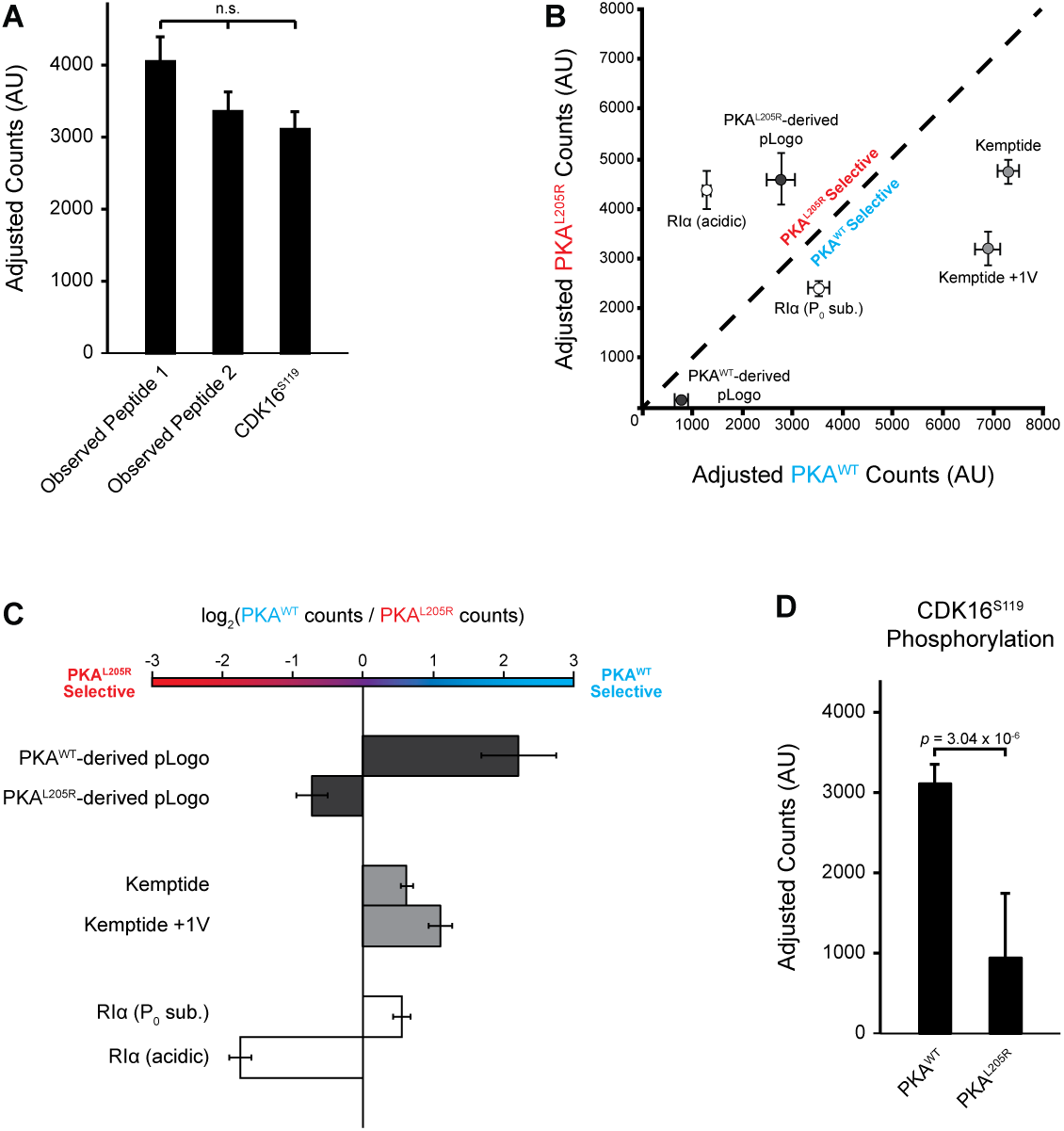
***In vitro* kinase assays confirm pLogo visualizations.** (A) Peptide arrays were used to evaluate PKA^WT^ phosphorylation of different candidate peptides by incubating membranes with recombinant kinase and [γ-^32^P]ATP in an *in vitro* kinase assay. Bars indicate adjusted counts (see Table S3) across technical replicates (*n* = 6-8), and error bars are ±SEM. A One-way ANOVA found the means are not significantly different [*F*(2,14) = 0.56, *p* = 0.58349]. (B) Peptide arrays were used to evaluate the phosphorylation activity of either PKA^WT^ or PKA^L205R^. Data points are color coded to indicate related candidate peptide pairs: pLogo-derived peptides (dark grey), Kemptide variants (light grey), and RIα variants (white). The dashed line indicates no preference (i.e. peptide is phosphorylated with equal efficiency by both PKA^WT^ and PKA^L205R^). Candidate peptides that fall below the line are phosphorylated at a higher efficiency by PKA^WT^, while candidate peptides that fall above the line are phosphorylated at a higher efficiency by PKA^L205R^. *n* = 6-8 technical replicates, and error bars are ±SEM. (C) Bars indicate the log_2_ ratio of PKA^WT^ activity to PKA^L205R^ activity. Candidate peptides that are phosphorylated at a higher efficiency by PKA^WT^ extend to the right, while candidate peptides that are phosphorylated at a higher efficiency by PKA^L205R^ extend to the left, and x = 0 indicates no preference. Error bars represent the propagated ±SEM. Peptide pairs are color coded as in the previous panel. (D) Phosphorylation of the known human PKA^WT^ substrate CDK16^S119^ is significantly reduced for PKA^L205R^ compared with PKA^WT^ (*n* = 8, t-test, p = 3.04 x 10^−6^). Error bars are ±SEM. Full blots and quantification are available in the Supporting information (Fig. S4 and Table S3). Peptides are: PKA^WT^-derived pLogo - RKRRRRKSFIENDRR, PKA^L205R^-derived pLogo - KRRRRRGSLDEDDQG, Kemptide - LRRASLG, Kemptide +1V - LRRASVG, Observed Peptide 1 - PLRMRRGSIPALVNN, Observed Peptide 2 - IVLPRRLSDEVADRV, RIα (P_0_ sub.) - KGRRRRGSISAEVYT, RIα (acidic) - KGRRRRGSDEAEVYT, CDK16^S119^ - EDINKRLSLPADIRL.

Next, we validated the observed specificity drift of the PKA^L205R^ mutation. We designed peptides based on the best residues at each position for each of the PKA^WT^ and PKA^L205R^ pLogos (RKRRRRKSFIENDRR and KRRRRRGSLDEDDQG, respectively) to assess the differential activity of each PKA variant for consensus substrates. While both peptides are viable substrates for both PKA variants, it is clear when comparing phosphorylation levels that the PKA^WT^ pLogo-derived peptide is preferentially phosphorylated by PKA^WT^, while the PKA^L205R^ pLogo-derived peptide is preferentially phosphorylated by PKA^L205R^ (Fig. 3B,C). This provides independent validation of the kinase specificity motifs identified by ProPeL using an orthogonal approach.

Having demonstrated that there is indeed a shift in substrate specificity, we sought to use this information to guide the design of peptide substrates with altered kinase selectivity. The gold-standard PKA consensus substrate Kemptide (LRRASLG) has been previously used to evaluate not only PKA^WT^ activity, but also PKA^L205R^ activity [5]. As mentioned above, PKA^WT^ displays a general preference for all hydrophobic residues in the P+1 position, while PKA^L205R^ retains only leucine from this hydrophobic grouping. As Kemptide contains arginine residues in the P-2 and P-3 positions and leucine at P+1, it conforms to our specificity models for both PKA variants. We hypothesized that replacing the Kemptide P+1 leucine with a valine (i.e. LRRASVG) would reduce phosphorylation by PKA^L205R^, while maintaining high phosphorylation levels by PKA^WT^. Indeed, our “Kemptide +1V” peptide exhibits increased selectivity towards PKA^WT^ over PKA^L205R^ when compared with standard Kemptide (Fig. 3B,C). Next, we investigated the biologically relevant regulatory subunit RIα interaction site (RRGAI). We hypothesized that by rationally mutating the RIα pseudo-substrate towards our model of PKA^L205R^ specificity, we could “rescue” the interaction between the Cushing’s mutant kinase and the RIα binding site. We synthesized an alanine-to-serine phosphoacceptor-substituted RIα peptide “RIα (P_0_ sub.)” to convert the interaction site to a potential substrate (KGRRRRGSISAEVYT). Based on our PKA^L205R^ ProPeL experimental findings, we also synthesized a version of the peptide with the putative +1 and +2 residues mutated to aspartate and glutamate, respectively, yielding an “RIα (acidic)” peptide (KGRRRRGSDEAEVYT). The *in vitro* kinase assays revealed that RIα (P_0_ sub.) is phosphorylated at a higher level by PKA^WT^, and that RIα (acidic) is phosphorylated at a higher level by PKA^L205R^ (Fig. 3B,C).

Finally, we wanted to test the feasibility of the hypothesis that the L205R mutation could induce substrate rewiring. Using the internal version of *scan-x*, we scored all of the known human PKA substrates (curated from the PhosphositePlus database [24]) with the PKA^WT^ and PKA^L205R^ motifs, and determined CDK16^S119^ as a candidate loss-of-function substrate for PKA^L205R^. The *in vitro* kinase assay revealed that phosphorylation of the CDK16^S119^ peptide by PKA^L205R^ was significantly reduced compared to PKA^WT^ (Fig. 3D, *n* = 8 technical replicates, *t*-test, *p* = 3.04 × 10^−6^). This result demonstrates both the feasibility of PKA^L205R^ loss-of-function mutations, and that ProPeL-generated specificity models can be utilized for predicting kinase substrate rewiring.

## Discussion

At present, disease-associated kinase mutations are largely classified as either inactivating or hyperactivating, and the potential impact of missense mutation on kinase specificity has not been fully explored [25,26]. Although several groups have noted an increase in PKA^L205R^ activity compared to wild-type, this activity has been assessed on only a small subset of known canonical PKA substrates (specifically CREB^S133^, ATF1^S63^, Kemptide and the PKA sensor AKAR4-NES) and in the presence of a regulatory subunit, typically RIα [4–7]. Therefore, it is important to distinguish whether an increase in phosphorylation of downstream PKA targets results from an actual increase in intrinsic PKA^L205R^ catalytic subunit activity, or is the result of constitutive PKA activity due to abolished regulatory subunit binding. Indeed, Lee et al. recently demonstrated a *reduction* in isolated PKA^L205R^ catalytic subunit activity [27], in agreement with our data (Fig. S2B,C). Our results, combined with previous findings, suggest that we should report the specific relationship of kinase activity with respect to individual substrates, rather than just reporting the activity on arbitrary substrates (which may not be universal). For example, when using the canonical Kemptide substrate, there is a clear *decrease* in activity of the isolated PKA^L205R^ catalytic subunit relative to PKA^WT^. However, our RIα (acidic) peptide shows the opposite result, with a clear *increase* in activity of PKA^L205R^ over PKA^WT^ (Fig. 3B,C, and Fig. S4). Both substrate selection and the absence of regulatory inhibition are therefore critical when designing and interpreting the results of a kinase catalytic activity assay. According to our model, many canonical substrates that conform to the PKA^WT^ [R/K][R/K]×Sϕ motif would remain viable targets for PKA^L205R^, in agreement with recent findings [4–7,11]. In fact, CREB^S133^ and ATF1^S63^ have identical substrate sites containing RRPSY, where we note that tyrosine is neither favored nor disfavored at the P+1 position. Because this peptide sequence matches favorably with both the PKA^WT^ and PKA^L205R^ motifs, these substrates would actually be *poor* candidates to evaluate a change in PKA^L205R^ specificity, and may explain why prior studies might have missed the altered specificity that we observe.

Altered PKA^L205R^ specificity suggests the possibility that gain-of-function PKA signaling might extend beyond hyperphosphorylation of canonical substrates to include phosphorylation of novel PKA substrates, and the loss-of-function of individual substrates. While it is beyond the scope of this study to identify and validate PKA substrate rewiring in Cushing’s Syndrome, we demonstrated the feasibility of loss-of-function by showing that PKA^L205R^ exhibits significantly reduced activity towards one example PKA substrate, CDK16^S119^, and gain-of-function with increased activity towards non-canonical peptide substrates. Therefore, while the model that PKA^L205R^ exhibits constitutive activity due to abolished regulatory subunit binding remains convincing, it is possible that the L205R mutation has additional effects resulting from an alteration in PKA’s pool of target substrates.

In this study, we have demonstrated that the Cushing’s Syndrome PKA^L205R^ mutation exhibits altered PKA substrate specificity. Using [γ-^32^P]ATP *in vitro* kinase assays we validated this specificity change, utilized our experimentally determined kinase motifs to engineer PKA variant-selective substrates, and demonstrated the feasibility of PKA^L205R^ loss-of-function. Together, our results suggest that the L205R mutation may influence Cushing’s Syndrome disease progression beyond simple hyperactivation of canonical PKA signaling, and demonstrate a powerful approach for investigating kinase substrate rewiring in human disease.

## Acknowledgements

The authors wish to thank Noah Dephoure and Craig Braun in the Gygi Lab for their assistance and support with mass spectrometry experiments. We thank Sean Lubner, John Redden, Stephanie Reeve, Megha Sah, Ben Stranges, and Randall Walikonis for generously sharing reagents and/or providing invaluable technical advice. We also thank Prakhar Bansal for maintainance and assistance with the Schwartz Lab computational tools. We thank Sean Congdon, Lylah Deady, Fred Murphy, Benjamin Currall and Anastasios Tzingounis for helpful discussions and the critical reading of our manuscript. Finally, we thank the University of Connecticut Computational Biology Core for hosting the pLogo web site, and maintaining the clusters on which it runs. This work was supported in whole or in part by grants awarded to GMC from the Department of Energy (DE-FG02-02ER63445), and to DS from the University of Connecticut Research Foundation, the University of Connecticut Office of the Vice President for Research, and the National Institute of Neurological Disorders and Stroke (1R21NS096516).

## Author Contributions

JML and DS conceived of the study. JML, MFC, and DS designed the experiments. JML, KLD-K, and MFC performed the experiments and analyzed the data. CRC, GMC and DS contributed materials, resources, and analysis tools. JML wrote the manuscript. All authors helped edit the final manuscript.

